# Bacterial RNA promotes proteostasis through inter-tissue communication in *C. elegans*

**DOI:** 10.1101/2024.03.13.584467

**Authors:** Emmanouil Kyriakakis, Chiara Medde, Danilo Ritz, Geoffrey Fucile, Alexander Schmidt, Anne Spang

## Abstract

Life expectancy has been increasing over the last decades, which is not matched by an increase in healthspan. Besides genetic composition, environmental and nutritional factors influence both health- and lifespan. Diet is thought to be a major factor for healthy ageing. Here, we show that dietary RNA species extend healthspan in *C. elegans*. Inherent bacterial-derived double stranded RNA reduces protein aggregation in a *C. elegans* muscle proteostasis model. This beneficial effect depends on low levels of systemic selective autophagy, the RNAi machinery in the germline, even when the RNA is delivered through ingestion in the intestine and the integrity of muscle cells. Our data suggest a requirement of inter-organ communication between the intestine, the germline and muscles. Our results demonstrate that bacterial-derived RNAs elicit a systemic response in *C. elegans*, which protects the animal from protein aggregation during ageing. We provide evidence that low stress levels are beneficial for healthspan.

**One-Sentence Summary:** Bacteria-derived dietary cues and inter-tissue communication promote proteostasis and fitness in *C. elegans*

## Introduction

Humans and all living organisms rely on nutrients for growth, reproduction, movement and survival, with key nutritional pathways being evolutionary conserved across species. It is generally accepted that the type and concentration of nutrients influence healthspan and life expectancy of eukaryotes. However, it remains unclear what combination of nutrients is most beneficial. Over the years, *C. elegans* has been proven to be an important and reliable model for nutrient-dependent health- and lifespan studies with major discoveries being confirmed across species (*1–5*). The influence of dietary restriction on longevity was first assessed in *C. elegans* and is now widely accepted for mammals and even humans (*3*, *6–8*). Furthermore, pioneering studies in *C. elegans* have unveiled the important role of cellular protein homeostasis (or proteostasis) in diseases and ageing (*9–14*). Since proteostasis deteriorates during ageing, finding ways to safeguard or even extend proteostasis emerges as a key concept to prevent, or at least ameliorate, age-associated diseases, such as cardiovascular disease, neurodegenerative diseases, late-onset neuromuscular disorders, sarcopenia and others. Ample scientific evidence suggests that specific dietary interventions is a promising approach to maintain proteostasis and improve health during ageing.

To answer how diet and which dietary components influence cellular and organismal fitness and life expectancy in a reliable and expeditious way, we investigated *C. elegans* and its bacterial diet. *C. elegans* nematodes are reared on monoxenic bacterial cultures that are easy to grow and to genetically manipulate. Utilizing this simple, tractable animal model, we show that a mixed diet of two *E. coli* strains promotes *C. elegans* fitness. Importantly, we demonstrate that bacterially-expressed ribonuclease 3 influences the accumulation of protein aggregates in *C. elegans* body-wall muscles, *via* a cell-non autonomous mechanism involving intestinal uptake of bacterial-derived RNA species, the RNAi machinery, selective autophagy and proper muscle function. We also show that communication across tissues and cell types, such as intestine, germline and neurons plays an important role in the regulation of proteostasis in body-wall muscles. Overall, our findings suggest bacterial-derived dietary cues influence organismal fitness by eliciting a protective response during stress and reveal how diet-derived RNA species promote proteostasis in *C. elegans*.

## Results

### A mixed bacterial diet promotes *C. elegans* fitness

In the laboratory, *C. elegans* are usually reared on two different *E. coli* strains: OP50, an *E. coli* B strain, and HT115, a K-12 derived strain. It was previously shown that these strains differ in their metabolic and nutrient profile (*15*). For example, OP50 leads to vitamin B12 deficiency in *C. elegans* (*16*, *17*). We first investigated the effect of diet on organismal fitness and lifespan (Fig. 1A). As previously reported, we did not observe any considerable difference in lifespan between the OP50 and HT115 diets (Fig. 1B) (*15*). However, worms fed on OP50 produced significantly higher number of progeny, which also developed faster than worms on the HT115 diet (Fig. 1C, D & Fig. S1). This beneficial effect came at the cost of reduced healthspan at advanced age, since a larger fraction of OP50-fed worms displayed impaired movement compared to their HT115-fed counterparts (Fig. 1E). These data indicate there were apparent benefits and trade-offs accompanying each diet. Hence, we reasoned that a mixture of both diets could exert beneficial effects. Indeed, the benefits of OP50 were still maintained even if it constituted only 10% of the diet, while the fitness in older worms was improved even beyond the level of feeding on HT115 alone (Fig. 1C-E, movies 1-4). Thus, bacterial diets differentially affect development, reproduction and healthspan. Combining both diets also combined the benefits of each individual diet and improved the healthspan of *C. elegans*.

**Figure 1.**
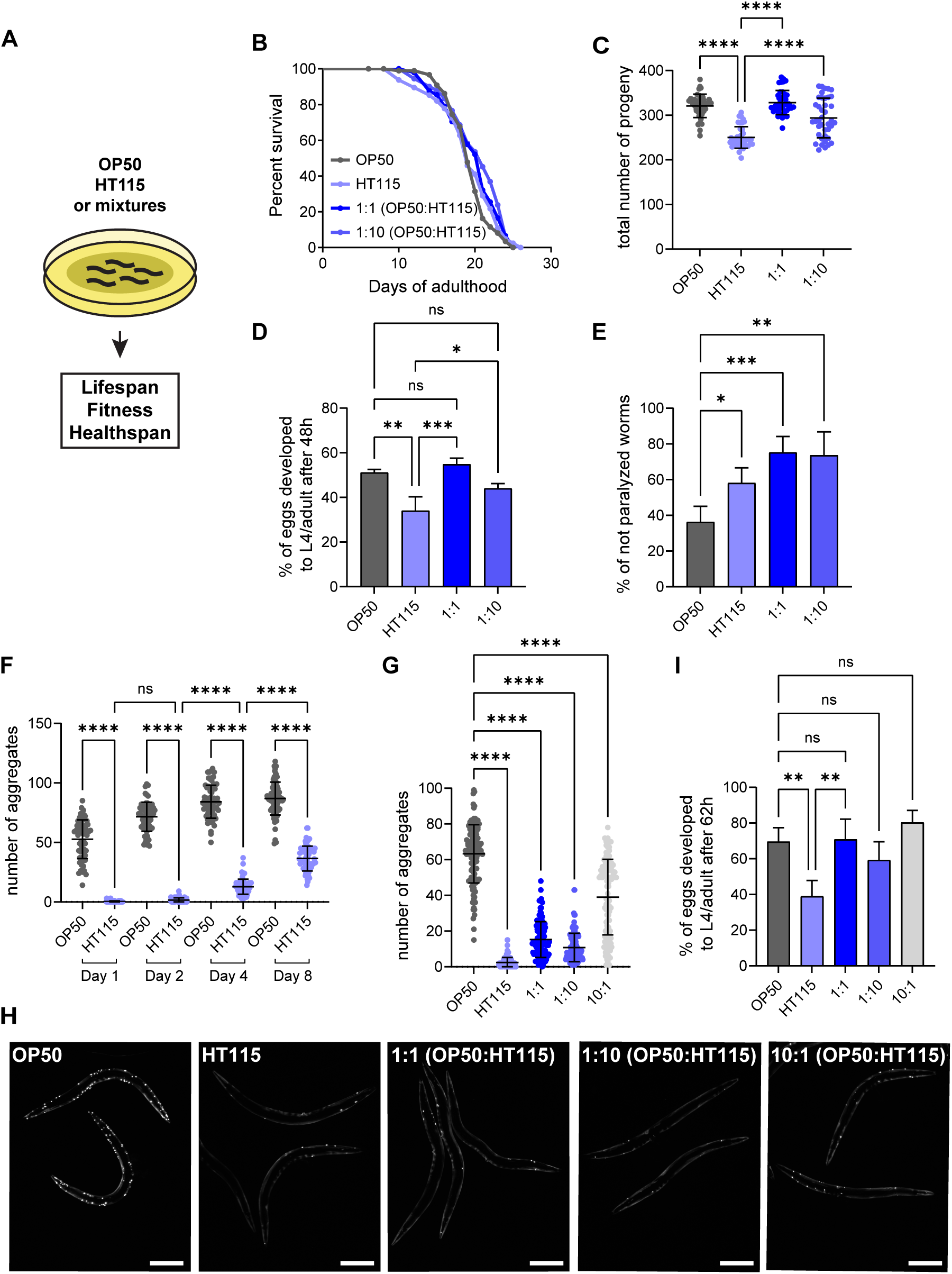
Dietary mixtures promote fitness and proteostasis in the body wall muscles of *C. elegans*. (A) Schematic representation of main methodology. Worms were grown on different bacteria lawns. Lifespan, fitness and healthspan assays where performed. (B) Lifespan curves of wt worms cultured on OP50, HT115 and bacterial mixtures. Bacterial mixtures of OP50 and HT115 were used in 1:1 (50% OP50 and 50% HT115) and 1:10 (10% OP50 and 90% HT115) ratios. Statistical analysis for lifespan curves was performed with the Log-rank (Mantel-Cox) test and the summary is shown in Table 1. (C) Brood size of wt worms on different bacterial lawns. The total number of progeny per worm is shown. (D) Developmental rate of wt worms on different bacterial lawns was measured as the percentage of eggs that developed into L4/adult stages 48h after egg laying. (E) Percentage of not paralyzed (18-20 days old) wt worms on different bacterial diets was assessed. Values represent mean ± SD from three independent experiments. (F) Quantification of polyQ40::YFP fluorescent foci of worms on OP50 or HT115 diet during ageing. The number of aggregates per worm is shown. (G) Quantification of polyQ40::YFP fluores-cent foci of 2-day old worms on different bacterial diets. The number of aggregates per worm is shown. (H) Representative images of 2-day old polyQ40::YFP-expressing worms on different bacterial diets. Scale bars in all panels are 200μm. (I) Developmental rate of polyQ40 expressing worms on different bacterial lawns was measured as the percentage of eggs that developed into L4/adult stages 62h after egg laying. Values represent mean ± SD from at least three independent experiments. One-way ANOVA with multiple comparison test was used. ns P>0.05, * P<0.05, ** P<0.01, *** P<0.001, **** P<0.0001.

### Dietary cues protect from muscle proteotoxicity

Next, we aimed to uncover how diets affect fitness and healthspan. Motility is a fitness measure in *C. elegans* and is linked to the function of body-wall muscles. Polyglutamine (polyQ) expansions have been used to assess cellular dysfunction in *C. elegans* body-wall muscles in response to proteotoxicity (*14*). polyQ40-YFP aggregate formation in body-wall muscles of worms can be used as a readout for proteostasis decline. Worms showed numerous polyQ-aggregates early in adulthood when reared on OP50, the number of which increased with age (Fig. 1F). In contrast, HT115-fed worms started to form aggregates much later in adulthood and to a lesser extent (Fig. 1F). The strong delay in aggregate formation in body-wall muscle cells might serve as indication for the increased mobility of aged animals on HT115 diet. These differences were not due to OP50’s vitamin B12 deficiency (Fig. S2). As expected, polyQ24-expressing worms did not show any diet-dependent aggregate formation (Fig. S3A). To corroborate the diet-dependent effect on proteostasis, we expressed amyloid-beta (Aβ1-42) in body-wall muscle cells, which has been previously shown to lead to paralysis in worms (*18*). Again, HT115 led to a much later onset of paralysis compared to OP50 (Fig. S3B). Mixed diets also improved the fitness of polyQ40-expressing animals, as we observed beneficial effects on the number of aggregates, number of progeny and development (Fig. 1G-H & Fig. S4A, B). Thus, we detected diet-dependent proteostasis dysregulation in body-wall muscles, and for which the polyQ40 *C. elegans* can serve as a sensor.

### Autophagy protects from diet-dependent accumulation of protein aggregates

One explanation for the positive dietary effect of the HT115 bacterial diet could be the upregulation of cytoprotective mechanisms, such as autophagy. Autophagy positively influences health- and lifespan by removing damaged organelles and protein aggregates (*19*). Moreover, it has been shown that autophagy inhibits the accumulation of polyQ40 aggregates in *C. elegans* and protects from proteotoxicity (*20*). Similarly, we found that knockdown of key autophagy factors increased the accumulation of polyQ40 aggregates on HT115 diet (Fig. 2A, B). Moreover, the offspring of mothers with defective autophagy were developmentally arrested (Fig S5A), and this effect was dependent on the expression of polyQ40 in muscle cells (Fig. S5B-D). It is plausible that the proteotoxicity of polyQ40 in muscle cells triggers a signal under these conditions, which is inherited by the offspring. Moreover, these data indicate that HT115 might induce autophagy systemically, which is beneficial for the animal. To test this possibility, we measured autophagy induction using *C. elegans* LC3 fused to GFP (LGG-1::GFP) in hypodermal seam cells as read-out for systemic induction. Autophagy was moderately but significantly induced by the HT115 diet compared to OP50 as measured by the number of autophagosomes present in hypodermal seam cells (Fig 2C, D). Protein aggregates are usually removed through a selective autophagy pathway termed aggrephagy. To test whether HT115 might induce aggrephagy, we knocked down E3 ligases implicated in the ubiquitination of aggrephagy clients, PELI-1 and CHN-1, the aggrephagy receptors SQST-1 (p62) and TLI-1 (Tollip/Cue) and the aggrephagy adaptor WDFY-3 (Alfy) (*21*, *22*). In all cases, we observed an increase in polyQ40 aggregate formation (Fig 2E, F & Fig. S5E, F), indicating that aggrephagy in *C. elegans* body-wall muscle cells is activated to prevent aggregate formation. Similar results were obtained in a *tli-1* KO strain (Fig. S5G, H). TLI-1 and SQST-1 appear to have at least partially overlapping functions as the combined knockdown increased aggregate formation over the individual knockdowns (Fig. 2G). Our data so far indicate that the HT115 bacterial diet protects from protein aggregation by inducing systemic autophagy, and that the OP50 diet cannot induce the response to similar levels.

**Figure 2.**
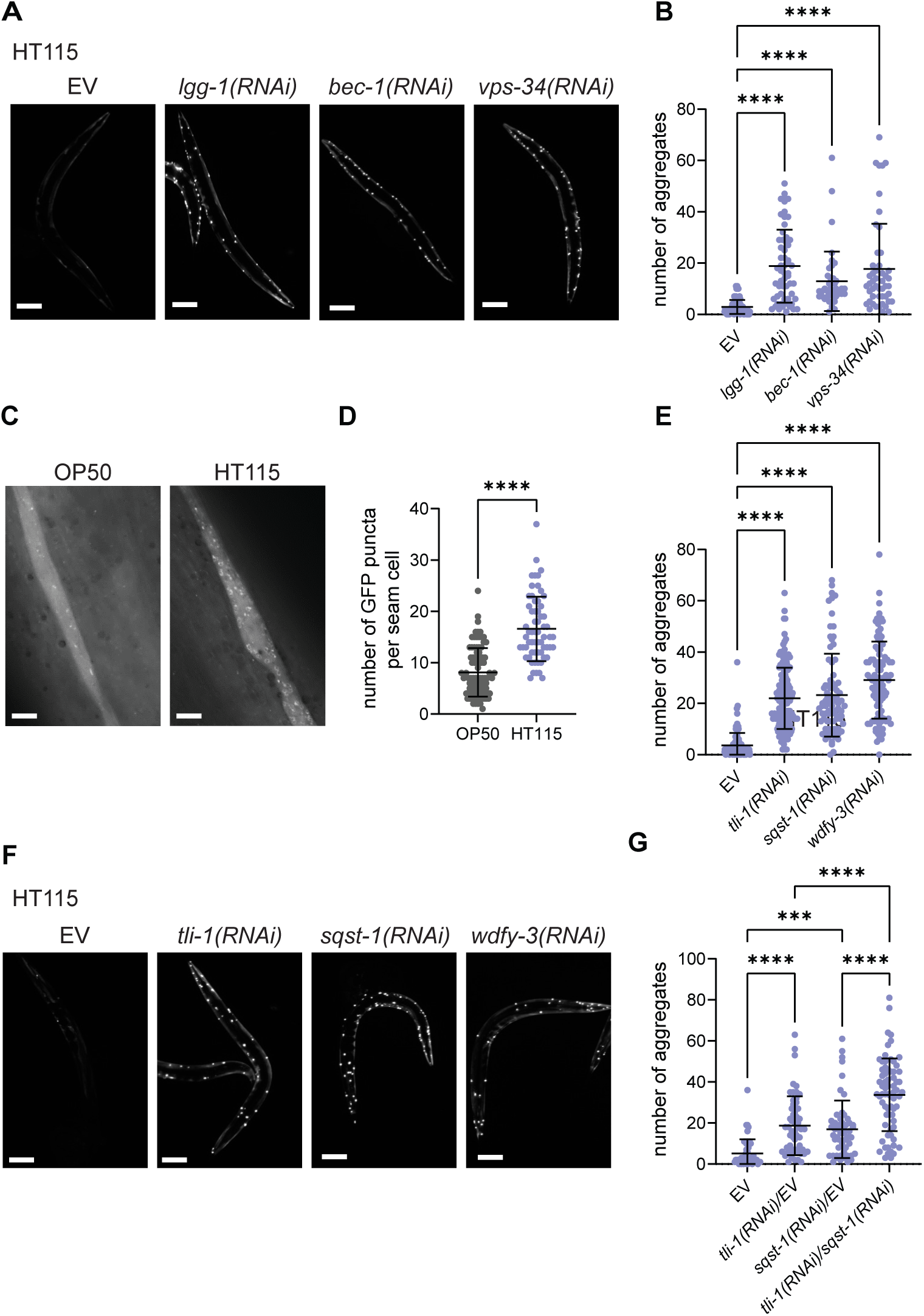
Autophagy protects from protein aggregation in the body wall muscles of *C. elegans*. (A) Representative images of 2-day old polyQ40::YFP-expressing worms on HT115 diet, treated with empty vector (EV), *lgg-1(RNAi), bec-1(RNAi) and vps-34(RNAi)*. (B) Quantification of polyQ40::YFP fluorescent foci of 2-day old worms on HT115, treated with empty vector (EV), *lgg-1(RNAi), bec-1(RNAi)* and *vps-34(RNAi)*. The number of aggregates per worm is shown. (C) Representative images of hypothermal seam cells, of p_lgg-1_GFP::LGG-1-expressing L4 animals on OP50 or HT115. Scale bar is 10μm (D) Quantification of GFP::LGG-1 positive puncta per seam cell of worms on OP50 and HT115. (E) Quantification of polyQ40::YFP fluorescent foci of 2-day old worm on HT115, treated EV, *tli-1(RNAi), sqst-1(RNAi)* and *wdfy-3(RNAi)*. The number of aggregates per worm is shown. (F) Representative images of 2-day old polyQ40::YFP-expressing worms on HT115 diet, treated with EV, *tli-1(RNAi), sqst-1(RNAi)* and *wdfy-3(RNAi)*. (G) Quantification of polyQ40::YFP fluorescent foci of 2-day old worms treated with EV, *tli-1(RNAi)* and *sqst-1(RNAi)* diluted with equal amount of the EV and mixture of equal amounts of *tli-1(RNAi)* and *sqst-1(RNAi)*. The number of aggregates per worm is shown.

### Innate immunity pathways are not activated by OP50 or HT115 diets

The positive dietary effect on proteostasis could also be due to stimulation of the innate immune response in worms. Induction of innate immunity has been observed with live pathogenic bacteria (*23*, *24*). To test this possibility, we fed polyQ40 worms with UV-killed OP50 and HT115. However, the diet-dependent aggregate formation remained unchanged (Fig. S6A, B). We next tested whether a bacterial secreted factor could be responsible for the differences between OP50 and HT115 and would induce innate immunity, which was not the case as the secretome of the bacteria had no beneficial effect on the worm proteostasis (Fig. S6C, D). Finally, we tested the induction of innate immunity more directly with transcriptional reporters for several immunity response genes. While those genes were induced by a pathogenic strain of *Pseudomonas aeruginosa* (PA14), neither OP50 nor HT115 elicited a response under the conditions tested (Fig. S7A-C). Thus, it is unlikely that the innate immune response is a prominent driver of diet-dependent aggregate formation.

### Ribonuclease-dependent bacterial-RNA species promote proteostasis

A key difference between OP50 and HT115, besides OP50 being a vitamin B12 auxotroph, is that HT115 can be used for RNA silencing experiments by feeding, while OP50 cannot. HT115 lacks a functional ribonuclease 3, which recognizes dsRNA species and cleaves them with high specificity to produce smaller dsRNA fragments. Therefore, ectopically expressed dsRNA in HT115 is stable and can be transferred to worms when they feed on the bacteria. To explore whether the presence or absence of ribonuclease 3, is important for the dietary difference in proteostasis in worms, we reared worms on the OP50(xu363) strain, in which *rnC* gene is mutated (*25*). Strikingly, loss of *rnC* protected worms from polyQ40 aggregates (Fig. 3A, B). Re-introduction of wild-type *rnC,* but not of two different catalytically inactive mutant versions (*26*, *27*) into OP50(xu363) or HT115 led to aggregate formation in animals, confirming the detrimental effect of bacterial *rnC* on worm proteostasis (Fig. 3C, D, Fig. S8). The HT115 parental strain (W3310), which still carries a functional *rnC* gene, caused aggregate formation in muscle cells, validating that indeed the loss of bacterial *rnC* improves *C. elegans* proteostasis (Fig. S9). Finally, we wanted to test the general applicability of our findings. The *E. coli* strain Nissle 1917 is a non-pathogenic bacterial strain used in the clinical setting to treat gastrointestinal conditions due to its probiotic properties. We deleted *rnC* in Nissle 1917. PolyQ40 expressing worms on *rnC*-ablated Nissle 1917 diet showed a significantly reduced number of aggregates in their body-wall muscles compared to worms fed with WT Nissle 1917 (Fig. 3E, F). Taken together, our data establish that loss of bacterial ribonuclease 3 has a positive effect on proteostasis in *C. elegans* muscle cells.

**Figure 3.**
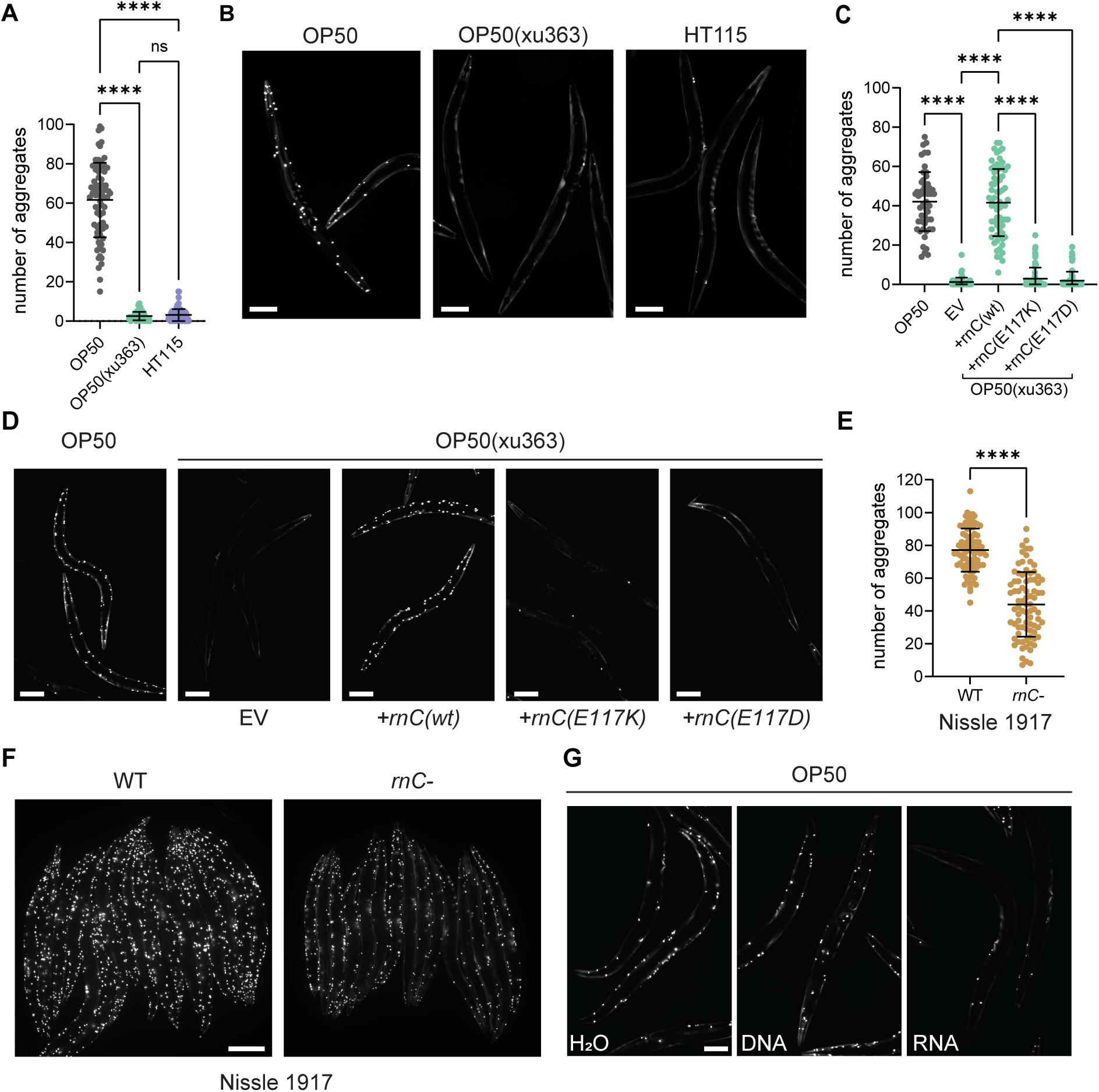
Ribonuclease 3-dependent bacterial-RNA species protect from polyQ40 protein aggregation. (A) Quantification of polyQ40::YFP fluorescent foci of 2-day old worms on OP50, OP50(xu363) and HT115 bacterial diets. The number of aggregates per worm is shown. Values represent mean ± SD from three independent experiments. One-way ANOVA with multiple comparison test was used. ns P>0.05, **** P<0.0001. (B) Representative images of 2-day old polyQ40::YFP-expressing worms on OP50, OP50(xu363) and HT115 bacterial diets. Scale bar is 100μm. (C) Quantification of total polyQ40::YFP fluorescent foci of 2-day old worms on OP50 and OP50(xu363) containing the empty vector (EV) or expressing the wt ribonuclease 3 (*+rnC(wt)*) and catalytically inactive ribonuclease 3 (*+rnC(E117K), +rnC(E117D)*). The number of aggregates per worm is shown. Values represent mean ± SD from three independent experiments. One-way ANOVA with multiple comparison test was used. **** P<0.0001. (D) Representative images of 2-day old polyQ40::YFP-expressing worms on OP50 and OP50(xu363) expressing the wt (*+rnC(wt)*) or catalytically dead (*+rnc(E117K)* or *+rnC(E117D)*) ribonuclease 3. EV serves as the control vector. Scale bar is 100μm. (E) Quantification of polyQ40::YFP fluorescent foci of 2-day old worms on wt and ribonuclease depleted (*rnC-*) Nissle 1917 E. coli. The number of aggregates per worm is shown. (F) Representative images of 2-day old polyQ40::YFP-expressing worms on wt and ribonuclease 3 depleted (*rnC-*) Nissle 1917 E. coli. Scale bar is 200μm. Values represent mean ± SD from three independent experiments. Student’s t-test was used. **** P<0.0001. (G) Representative images of 2-day old poly-Q40::YFP-expressing worms on OP50 diets, descendants of worms in which their gonads were injected with water (H2O), DNA or HT115-derived RNA. Scale bar is 100μm.

We confirmed that similar to HT115, the loss of aggregate formation on OP50(xu363) diet was also due to autophagy, and more specifically to aggrephagy (Fig. S10). Moreover, mutation of *rnC* in OP50 did not negatively influence brood-size or development (Fig. S11). Thus, the OP50(xu363) diet combines the advantages of the OP50 and HT115 diets with respect to organismal fitness. Our data suggest that bacterially-derived RNA species have a positive systemic effect on *C. elegans* proteostasis. To corroborate these findings, we injected RNA from bacteria or genomic DNA into the gonad of polyQ40 expressing *C. elegans.* Only bacteria-derived RNA reduced aggregate formation in muscle cells (Fig. 3G). These data suggest that loss of one single bacterial gene, *rnC,* provides essential RNA species through either the intestine or the germline that are critical for the reduction of aggregates in *C. elegans*.

### The RNAi machinery and the germline are required for the diet-dependent aggregate accumulation

*C. elegans*, like other eukaryotes, has evolved a system to defend itself against foreign RNA species, the RNAi machinery (*28*). Therefore, we tested whether bacterial-derived RNAs act through the RNAi machinery to promote proteostasis. To that end, we knocked-down key components of the RNAi pathway: the RNA transporters SID-1 and SID-2 and the Argonaute proteins RDE-1 and ERGO-1 using the HT115 diet. In all these cases, the number of aggregates in muscle cells was increased (Fig. 4A). A similar result was obtained upon silencing of the RNA-dependent RNA polymerase EGO-1, which is required for the systemic effect of RNAi in worms (Fig. 4A). These results were confirmed by mutants in RNAi pathway components (Fig. 4B-F), irrespective of the diet. Thus, our data provide strong evidence that bacterial-derived RNA is recognized by the RNAi machinery and that this machinery is linked to the reduction of polyQ40 aggregation.

**Figure 4.**
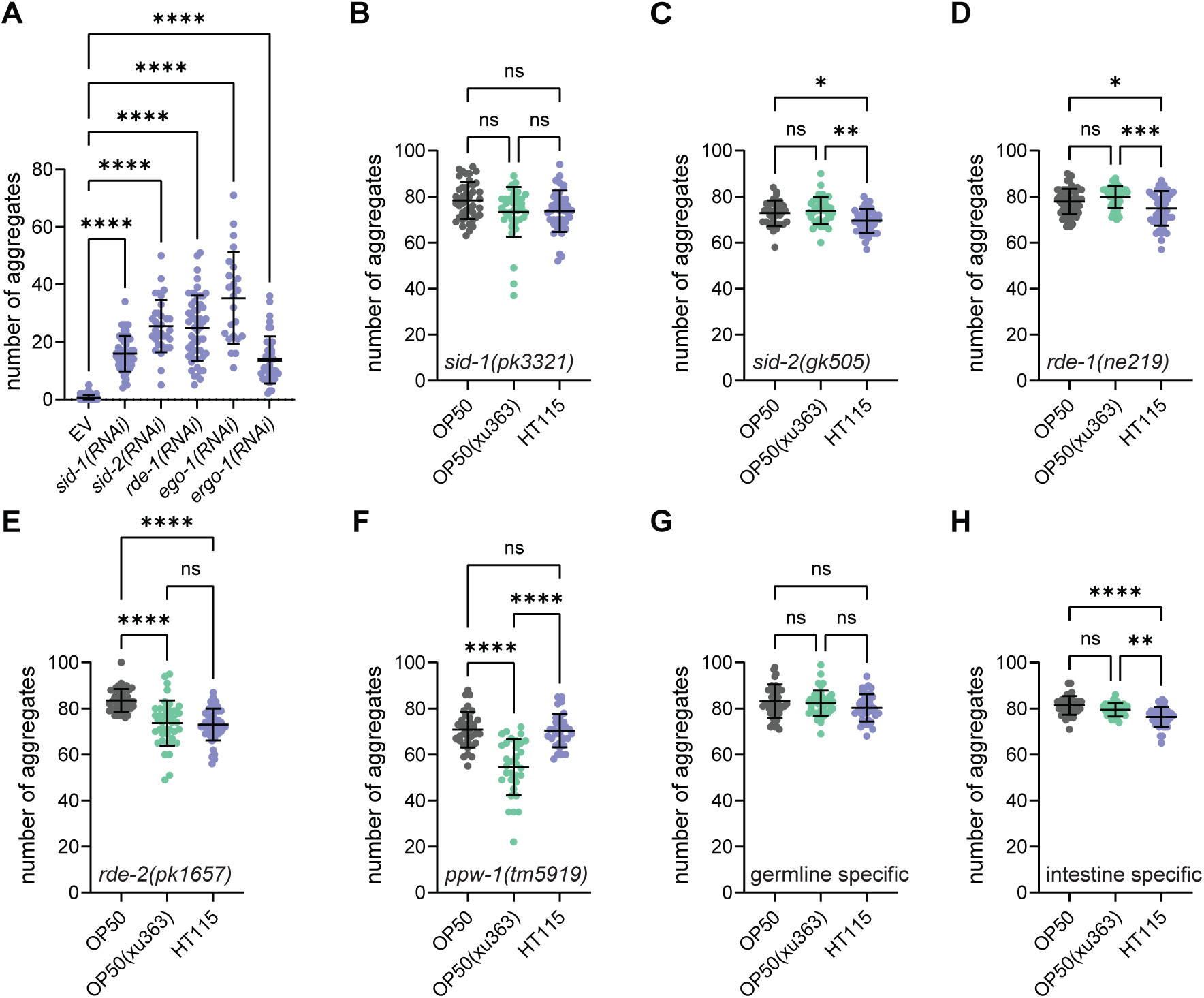
The RNAi machinery and the germline are required to protect from protein aggregation. Quantification of polyQ40::YFP fluorescent foci of 2-day old worms on HT115, treated with empty vector (EV), *sid-1(RNAi), sid-2(RNAi), rde-1(RNAi), ego-1(RNAi)* and *ergo-1(RNAi)*. The number of aggregates per worm is shown. (B-F) Quantification of polyQ40::YFP fluorescent foci of 2-day old worms on OP50, OP50(xu363) and HT115. The number of aggregates per worm is shown in *sid-1(pk3321)* (B), *sid-2(gk505)* (C), *rde-1(ne219)* (D), *rde-2(pk1657)* (E) and *ppw-1(tm5919)* (F) strains. (G) Germline and (H) intestine RNAi-specific mutant strains were used to quantify polyQ40::YFP fluorescent foci of 2-day old worms of OP50, OP50(xu363) and HT115. The number of aggregates per worm is shown. Values represent mean ± SD from three independent experiments. One-way ANOVA with multiple comparison test was used. ns P>0.05, * P<0.05, ** P<0.01, *** P<0.001, **** P<0.0001.

We have shown above that administration of bacterial RNA through the intestine by feeding and into the gonad by injection prevents aggregate formation in body wall muscles. Therefore, we wondered whether the RNAi machinery needed to be active in both tissues. PPW-1 is a germline-specific Argonaute (*29*). A mutant in PPW-1 accumulated polyQ40 aggregates in muscle cells, independent of the diet (Fig. 4F), indicating that even RNA delivery into intestine requires the germline RNAi components for the protective effect. However, the germline RNAi machinery was not sufficient, because a *C. elegans* strain in which RNAi is only active in the germline still accumulated polyQ40 aggregates (Fig. 4G). Consistent with these findings, polyQ40 aggregates also formed in animals, when the RNAi machinery was only active in the intestine (Fig. 4H). These results suggest that a functional germline is indispensable to block polyQ40 aggregate accumulation in the body-wall muscles of *C. elegans*. Moreover, communication across tissues involving intestine cell, the germline and muscle cells, is required for bacterially-derived RNA species to protect from protein aggregates.

Next, we determined whether any or a specific RNA would elicit the beneficial effect. If it was a specific one, this bacterial RNA would affect gene expression of a selective group of genes in the animal. We performed total RNA seq and smRNA seq on the three bacterial strains using both total RNA and a library enriched for small RNA molecules. On the total RNA level, HT115 and OP50 are distinct with the majority of the variance captured by inter-strain differences. Only a minor fraction of the total variance is captured by the contrast of OP50 versus OP50(xu363) (Fig. S12A). Among the set of 271 genes, which were consistently differentially expressed when contrasting HT115 versus OP50 and OP50(xu363) versus OP50 (Fig. S12 B), there is no significant enrichment of known Gene Ontology (GO) terms at any level. However, when considering the subset of 23 genes which are consistently up-regulated in HT115 and OP50_xu363 compared to OP50 there are enrichments in GO biological pathways, in particular related to RNA and protein metabolism (Fig S12C). RNA-Seq of the small RNA library similarly yielded few differences between the genotypes (Fig. S12D). We employed de novo transcriptome assembly to increase sensitivity, however among the expressed contigs we identified only three which were differentially expressed between OP50 and OP50(xu363) and mapped to the *C. elegans* reference genome (Table 2). Administration of dsRNA based on these candidates did not affect the phenotype (Fig S12E). Thus, it is unlikely that the bacterial RNA silences a gene or subset of genes specifically, but rather suggests that bacterial RNA elicit a low level of a more general stress response.

### Body-wall muscle contraction and sarcomere integrity alleviate accumulation of aggregates

Nevertheless, the dietary bacterial RNA prompted a systemic response and therefore we determined the proteome of WT and polyQ40 expressing worms reared on OP50, OP50(xu363) or HT115 diets. We detected diet-dependent changes in the proteomes of both WT and polyQ40 expressing worms, most importantly also between OP50 and OP50(xu363) (Fig. 5 A-F). We focused on the beneficial effect in our proteostasis model. In total, we found 194 proteins to be significantly altered in polyQ40-expressing worms grown on HT115 or OP50(xu363) compared to OP50 (Fig. S13A). GO term analysis revealed an enrichment of GO terms related to sarcomere organization and muscle function (Fig. S13B, Table 3). The functional protein association network uncovered 12 muscle-related proteins that were clustering (Fig. S13C). These proteins were not significantly altered in WT worms (Fig. 5A-F), indicating that the bacterial-derived RNA can elicit a context-dependent response and increase muscle function.

**Figure 5.**
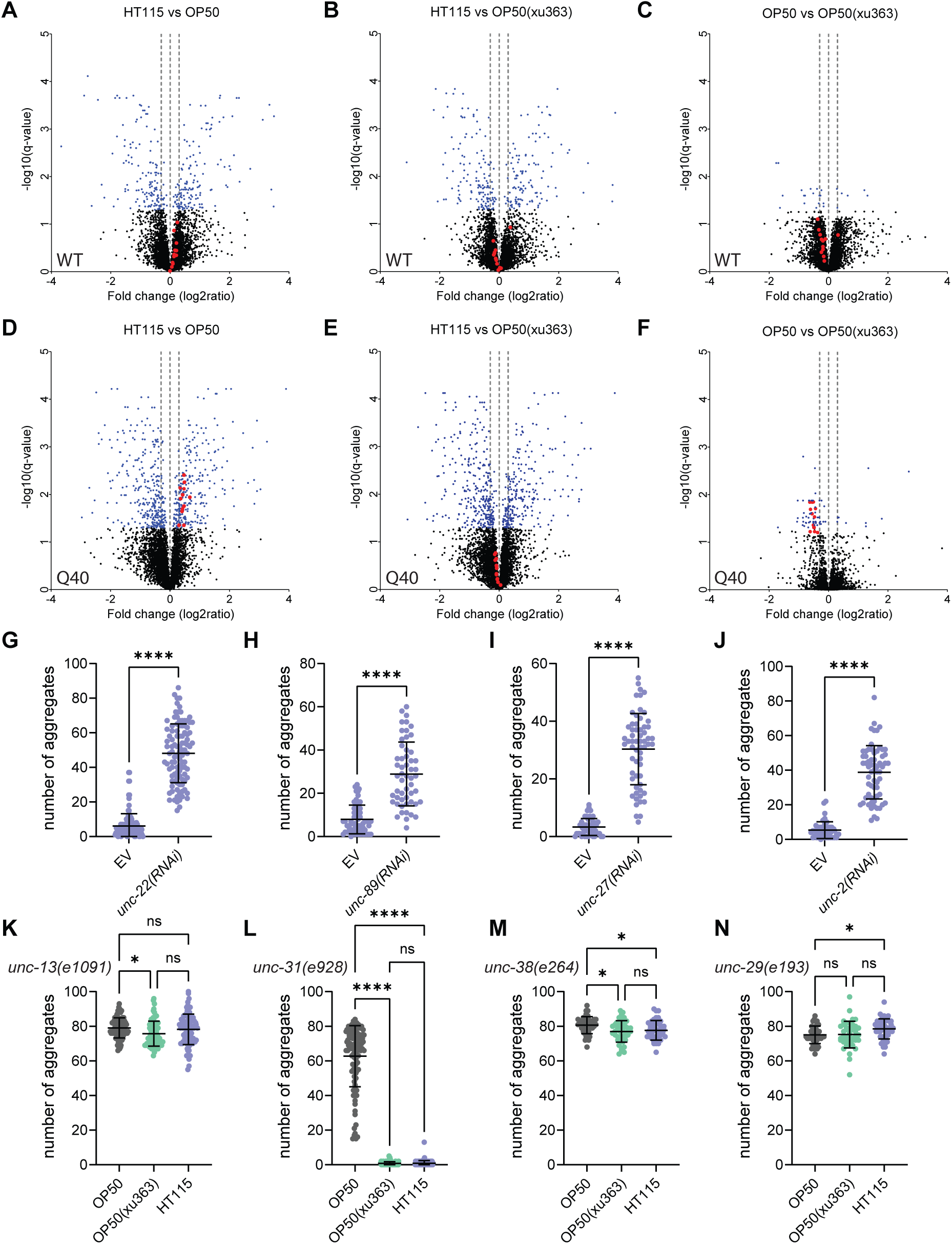
Muscle function and neurotransmission protects from accumulation of protein aggregates. (A-F) Volcano plots of total quantified proteins showing significant increase or decrease content in WT (A-C) and polyQ40-expressing (D-F) strains on OP50, OP50(xu363) or HT115 bacteria. In blue are the proteins with q values less than 0.05. UNC-89, UNC-22, TTN-1, UNC-15, ATN-1, UNC-54, UNC-87, ZK1321.4, Y43F8B.1, TNT-2, CPN-3, CLIK-1 proteins from the STRING analysis are shown in red. Horizontal dotted lines are at -0.3, 0, +0.3 fold change. (G-J) Quantification of polyQ40::YFP fluorescent foci of 2-day old worms on HT115, treated with empty vector (EV), *unc-22(RNAi)* (G), *unc-89(RNAi)* (H), *unc-27(RNAi)* (I) and *unc-2(RNAi)* (J). (K-N) Quantification of polyQ40::YFP fluorescent foci of 2-day old worms on OP50, OP50(xu363) and HT115. The number of aggregates per worm is shown in *unc-13(e1091), unc-31(e928), unc-38(e264)* (F) and *unc-29(e193)* (G) mutant strains. Values represent mean ± SD from three independent experiments. One-way ANOVA with multiple comparison test or Student’s t-test were used. ns P>0.05, * P<0.05, **** P<0.0001. Scale bars in all panels are 100μm.

To test this hypothesis, we performed knockout and knockdown experiments on key muscle genes. Lesion of two giant sarcomeric proteins, UNC-22 (twitchin) and UNC-89 (obscurin), or UNC-27 (the ortholog of troponin I) increased the number of protein aggregates (Fig. 5G-I & Fig. S14A-C). Sarcomere, the basic unit of muscles, contracts, following calcium influx. Disrupting calcium influx in the body-wall muscles by silencing *unc-2, egl-19* or *cca-1*, which encode subunits of three voltage-dependent calcium channels, increased the number of aggregates compared to the control (Fig. 5J & Fig. S14D-F). The voltage-dependent calcium channels open, when muscle membranes depolarize upon neuroendocrine stimulation. This stimulation could either happen through neurotransmitter release or neuropeptide secretion and processing (*30–32*). Mutants defective in the neurotransmitter release and acetylcholine synthesis (*unc-13* and *cha-1 cho-1*), but not in neuropeptide secretion and processing (*unc-31* and *egl-21*), lost the diet-dependent protective effect from polyQ40 aggregate formation in muscle cells (Fig. 5K, L & Fig. S15A-F). Likewise, blocking neurotransmission through mutation or silencing of the postsynaptic acetylcholine receptors UNC-38 and UNC-29 caused aggregate formation independent of the diet (Fig. 5M, N & S15G). In contrast, silencing acetylcholine receptors in motor or sensory neurons or the homomeric ACR-16 in body-wall muscles had no effect (Fig. S15H). These results provide evidence for the requirement of inter-tissue communication (Fig. S15I) to maintain functional body-wall muscles and protect from protein aggregation. Moreover, we find the dietary RNA mediates this protective effect by increasing the levels of key proteins required for muscle function.

## Discussion

Here, we provide evidence that atypical dietary components, in particular bacterial-derived RNA species regulate proteostasis and promote organismal health in *C. elegans*. Bacterial RNA species are taken up and processed by *C. elegans* intestinal cells and the germline to promote muscle function and protect from toxic protein aggregates *via* a mechanism that requires the RNAi pathway and autophagy induction. For the beneficial effect, inter-organ communication of the intestine, the germline and muscles is required (Fig. S16). Whether neuronal input acts in parallel or is also part of this communication system remains to be determined. These results suggest that loss of proteostasis is not solely an internal cellular problem but non-cell autonomous mechanisms are likewise important.

It has been shown previously that bacteria-derived RNAs can specifically affect gene expression of a subset of genes and behaviour in *C. elegans* (*33*, *34*). In a groundbreaking study, it was shown that pathogen-derived P11 sRNA is taken up by *C. elegans* and processed through the canonical RNAi pathway, to downregulate specifically the *maco-1* gene to initiate pathogen avoidance (*33*). The mechanism we uncovered in this study is different as we did not identify any specific bacterial RNA that would directly affect gene expression by acting as siRNA or by influencing transcription in *C. elegans.* These findings are mirrored by the proteome analysis in *C. elegans*, which did not reveal significant upregulation of major stress-responsive mechanisms nor a specific protein target. A major difference between our and previous studies is that we use a proteostasis model, which might already challenge the animals and therefore enabling them to better and faster adapt to stressful environments. Accordingly, our proteomic analyses revealed an increase in the concentration of proteins required for muscle function specifically in the muscle proteostasis model and not in the WT. Bacterial RNA(s) induce the expression of muscle specific genes to confer protection under stress conditions, suggesting that this trans-kingdom mechanism serves as a response, which is particularly beneficial during adverse and pathological conditions, indicating that organ specific responses could be triggered through dietary RNA and further supports the existence of broader, systemic effects. In support of this notion, we found systemic induction of selective autophagy. The relationship between proteostasis and autophagy is complex and context dependent. Moderate levels of autophagy are considered beneficial for cellular health. In our proteostasis model, autophagy is induced by the diet-derived RNA species and this is sufficient to promote proteostasis. Overall, the distinct response between control and proteotoxically challenged worms suggests an adaptive mechanism that depends on bacterial RNA and also the homeostatic state of cells and the whole organism. We propose a model in which the diet-derived RNA species would elicit a basal stress response that would prime the organism to deal better with the onset of protein aggregation in our proteostasis model and therefore reduces and delays aggregate formation in body-wall muscle cells. The stress response remains at a low level, which we assume is sufficient to trigger proteoprotective mechanisms on the cellular level and thereby reduce aggregate formation. This delay in aggregate formation may be the underlying cause of the increase in healthspan in our proteostasis model. The low level of stress induction is supported by our findings that we did not observe any significant upregulation of autophagic or other stress-response related proteins in our *C. elegans* proteomics analysis when the animals were fed the different bacterial diets.

It has been previously shown that diet may promote proteostasis and lifespan and protect from neurodegeneration in *C. elegans* (*1*, *2*, *35*, *36*). The transcriptional responses of *C. elegans* highly depends on the bacterial diet and different diets direct unique transcriptional signatures (*2*, *37*). In this study, we demonstrate that the transcriptional changes between the OP50 and HT115 diets are not responsible for the promotion of proteostasis and protection from toxic protein aggregates in *C. elegans*. However, the deletion of a single bacterial gene (*rnC*) in several distinct bacterial strains (OP50, HT115, Nissle1917) resulted in minimal transcriptional changes between the bacterial strains but significant changes in the *C. elegans* proteostasis model. Thus, the accumulation of dsRNA species that cannot be processed in the absence of ribonuclease 3 (*rnC*) is sufficient for proteoprotective effects observed in *C. elegans*.

In this study, we move beyond the strict definition of nutrients and we identified non-traditional components, such as RNA species, that promote proteostasis. Bacteria do not behave solely as a nutrient source and this interspecies model may be relevant in understanding the relationship between humans and their microbiome and how it impacts physiology and disease.

In particular, in the advent of RNA as therapeutics, it is conceivable that dietary small RNA will prove useful as intervention to extend healthspan in humans.

## Materials and methods

### Nematode strains and growth conditions

Standard rearing conditions were used for maintaining *C. elegans* strains. All experiments were performed at 20°C on nematode growth media (NGMs) agar supplemented with *Escherichia coli* (OP50, OP50(xu363), HT115, W3310, Nissle 1917, or mixtures of OP50 and HT115) unless otherwise stated. All bacteria strains were carrying the empty vector (EV) plasmid (pL4440) which served as the control for RNAi experiments but also for selection purposes, unless otherwise indicated. For RNAi experiments, worms were placed on NGM plates seeded with IPTG-induced HT115(DE3) or OP50(xu363) bacteria transformed with the gene-specific RNAi construct. OP50(xu363) is an OP50-derived RNAi-competent strain and HT115 is an RNAi-competent strain which derives from W3310 strain. For *egl-19(RNAi)* experiments, the bacteria cultures were diluted 10 times with the EV plasmid to minimize developmental defects and sterility. Clones of interest were obtained from the Ahringer RNAi bacterial library or generated in the lab. The following nematode strains were used in the study: N2: wild-type Bristol isolate, AM141: rmIs133 [punc-54Q40::YFP], AM138: rmIs130 [unc-54p::Q24::YFP], AM140: rmIs132 [unc-54p::Q35::YFP], CL4176: smg-1(cc546) I; dvIs27[myo-3p::A-Beta (1–42)::let-851 3’UTR) + rol-6(su1006)], MAH14: daf-2(e1370) III; adIs2122 [lgg-1::GFP + rol-6(su1006)], RB1473: tli-1(ok1724) (6 times outcrossed), VP303: rde-1(ne219) V; kbIs7 [nhx-2p::rde-1 + rol-6(su1006)], NR350: rde-1(ne219)V; kzIs20 [hlh-1p::rde-1 + sur-5p::NLS::GFP], KP2018: egl-21(n476) IV, DA509: unc-31(e928) IV, CB1091: unc-13(e1091) I, ppw-1(tm5919), DCL569: mkcSi13 [sun-1p::rde-1::sun-1 3’UTR + unc-119(+)] II; rde-1(mkc36) V, NL3321: sid-1(pk3321) V, WM27: rde-1(ne219) V, NL3531: rde-2(pk1657) I, VC1119: dyf-2&ZK520.2(gk505) III, CB193: unc-29(e193) I, CB904: unc-38(e264) I, RM1743: cha-1(md39) cho-1(tm373) IV, AY101: acIs101 [F35E12.5p::GFP + rol-6(su1006)], AU133: agIs17 [myo-2p::mCherry + irg-1p::GFP] IV, AU306: agIs44 [Pirg-4::GFP::unc-54–3′UTR; Pmyo-2::mCherry]. To generate double mutants, AM141 males were mated to hermaphrodites carrying the mutation of interest. The presence of the respective mutations was checked phenotypically or by genotyping.

### Constructs generated

For the construction of *tli-1(RNAi)* plasmid, the following primers were used: 5′-TCTAGAAACCAAAACAAATACTGATCTTCCGT-3’ (FW) (with XbaI restriction site) and 5′-ACCGGTCTCTCGGCTGCTGCTGTCATCT-3′ (RV) (with AgeI restriction site). The amplified *tli-1* genomic region was ligated into the pL4440 vector upon digestion with XbaI and AgeI. For the construction of *rnC*(wt) expression plasmid, the following primers were used: 5’-CCTGTGGATCCATGAACCCCATCGTAATTAATCG-3’ (FW) (with BamHI restriction site) and 5’-CCTGTCAGCTGTCATTCCAGCTCCAGTTTTTTC-3’ (RV) (with PvuII restriction site). The amplified *rnC* region was ligated into the pL4440 vector upon BamHI and PvuII digestion which leave only the one T7 promoter. For the construction of *rnC*(E117D) expression plasmid, the following primers were used: 5’-CCTGTGGATCCATGAACCCCATCGTAATTAATCG-3’ (FW) (with BamHI restriction site) and 5’-TAATGCATCGACGGTGTCGGCGA-3’ (RV) and also the 5’-CCTGTCAGCTGTCATTCCAGCTCCAGTTTTTTC-3’ (RV) (with PvuII restriction site) and 5’-TCGATGCATTAATTGGTGGCGTATT-3’. The two amplified regions were combined by fusion PCR using the following primers: CCTGTGGATCCATGAACCCCATCGTAATTAATCG-3’ (FW) (with BamHI restriction site) and 5’-CCTGTCAGCTGTCATTCCAGCTCCAGTTTTTTC-3’ (RV) (with PvuII restriction site). The amplified *rnC* region carrying the E117D point mutation was ligated into the pL4440 vector upon BamHI and PvuII digestion which leave only the one T7 promoter. For the construction of *rnC*(E117K) expression plasmid the same strategy was followed, the following primers were used: 5’-CCTGTGGATCCATGAACCCCATCGTAATTAATCG-3’ (FW) (with BamHI restriction site) and 5’-TAATGCTTTGACGGTGTCGGCGA-3’ (RV) and the 5’-CCTGTCAGCTGTCATTCCAGCTCCAGTTTTTTC-3’ (RV) (with PvuII restriction site) and 5’-TCAAAGCATTAATTGGTGGCGTATT-3’. For the construction of plasmids containing the sequences of the three expressed contigs which are differentially expressed between OP50 and OP50(xu363) that were identified from the RNA seq analysis, the following primers were used: (For hit 1) 5’-ctcgaattcACCCCATCGTAATTAATCGG-3’ (FW) and 5’-ctcgaattcTATTTTTAAAGTGATGATAAAAGGC-3’ (RV), (for hit 2) 5’-ctcgaattcTTTTAGCGTTTATATCTGAAGG-3’ (FW) and 5’-ctcgaattcCTTATGATGATGATGTGCTTAAA-3’ (RV), (for hit 3) 5’-ctggaattcTCAGCGCAATTGATAGGC-3’ (FW) and 5’-ctggaattcGTTTTTTCGCCCCATTTAG-3’ (FW), all containing EcoRI restriction site at the 5’. The amplified bacterial regions were ligated into the pL4440 vector upon digestion with EcoRI. These plasmids were used to generate dsRNA which were used to test their efficiency to modulate aggregate accumulation.

### Lifespan assays

Lifespan assays were performed at 20°C. Synchronous animal populations were generated by bleaching (hypochlorite treatment) gravid adult animals of the desired strain. Eggs were then placed on NGM plates with the different bacterial diets, until the L4 larval stage when they were again placed on the same diets. Their progeny was grown until the L4 larval stage and then transferred to fresh plates in groups of 20-25 worms per plate for a total of 100-120 individuals per condition (day 0 of adulthood). Animals were transferred to freshly-made RNAi plates every 2 days until the 12th day of adulthood and every 3 days until the end of the experiment. Animals were transferred to fresh plates every 2–3 days thereafter and examined every day for touch-provoked movement and pharyngeal pumping, until death. Worms that died owing to internally hatched eggs, an extruded gonad or desiccation due to crawling on the edge of the plates were censored and incorporated as such into the data set. Each survival assay was repeated at least twice and figures represent typical assays. Survival curves were created using the product-limit method of Kaplan and Meier.

### Brood size determination

Synchronous animal populations were generated by bleaching (hypochlorite treatment) of gravid adult animals. Eggs were then placed on NGM plates with the different bacterial diets and were grown on the same diet for at least two generations. Ten L4 worms were picked and placed into separate NGM plates containing the corresponding diet. After the first 36h worms were moved daily to fresh plates until no more eggs were laid. The number of progeny was scored in each plate and statistical analyses were performed using Sidak’s multiple comparisons tests following one-way ANOVA. Total or daily brood sizes are reported. Each brood size determination assay was repeated four times.

### Developmental rates

Synchronous animal populations were generated by bleaching (hypochlorite treatment) of gravid adult animals. Eggs were then placed on NGM plates with the different bacterial diets and were grown on the same diet for at least two generations. Each time, L4 worms were used to obtain synchronous worm populations. Approximately 12 2-day old worms were used for egg laying for 2-3h on fresh plates. The adult worms were removed and the number of eggs was determined. When approximately 50% of the worms started reaching the L4/adult stage, we counted the total number of progeny that reached the L4/adult stage (for different strains different time point was used due to developmental differences between strains). We performed statistical analyses using Sidak’s multiple comparisons tests following one-way ANOVA. The assay was repeated at least three times.

### Analysis of polyQ protein aggregation

For the analysis of polyglutamine aggregation in body wall muscle cells, we used the AM141 (polyQ40) and AM138 (polyQ24) strains. Synchronous animal populations were generated by bleaching (hypochlorite treatment) of gravid adult animals. Eggs were placed on NGM plates with the different bacterial diets and were grown on the same diet for at least two generations. L4 worms were used to obtain synchronous worm populations. Same age adult worms were collected, immobilized with levamisole before mounting on coverslips for microscopic examination with a Zeiss Axioplan 2 epifluorescence microscope. Protein aggregates in whole animals were quantified with the help of ImageJ software.

### Motility assay

Synchronous nematodes were grown normally on NGM media plates containing the different diets for at least 2 generations. When worms reached 17-20-days old, they were gently touched with an eyebrow. Worms not responding to touch were considered dead and were excluded from the analysis. Worms responding to touch but did not move were scored as paralysed and the percentage of paralysed worms per condition was evaluated. Each analysis was performed three times.

### Paralysis assay

To assay β-amyloid toxicity we used the CL4176 temperature sensitive strain that expresses β-amyloid in the body-wall muscle of *C. elegans,* leading to paralysis. Synchronous animal populations were generated by bleaching (hypochlorite treatment) gravid adult animals. Eggs were placed on NGM plates with the different bacterial diets at 15°C. At L4 stage 12 animals were transferred to fresh plates for two days. Then, egg laying was performed for approximately 3-4h. The mothers were removed and the plates containing only the eggs were placed back at 15°C for 48h, at which point the plates were shifted at 23°C. Approximately 24h later and for every 1-2h paralysis was scored. Percentage of paralyzed animals per conditions is plotted against the time since temperature shifting. Each analysis was repeated at least three times.

### Supernatant isolation and supplementation

Overnight OP50 and HT115 bacterial cultures (6 ml) were centrifuged for 10 min at 16, 000 rcf at 4°C. Bacterial supernatants were centrifuged for another 5 min and sterile-filtered using 0.2 μm filters. One ml of each supernatant (or LB medium as control) was used to overlay NGM media plates. The pellets were once washed with LB medium (5 ml), resuspended in 0.5 ml cold LB medium and spotted onto NGM media plates (100 μl). Plates were exposed to UV light for 20 min to kill the bacteria. Worms were reared on these plates for two generations and polyQ40::YFP protein aggregate were monitored.

### Methylcobalamin supplementation

The bacterial growth media was supplemented with exogenous methylcobalamin (Sigma), a vitamin B12 analog, to a final concentration of 25 μg/ml. OP50 bacteria were grown for two hours at 37 degrees, and spotted onto NGM media plates. Worms were reared on these plates for two generations and polyQ40::YFP protein aggregates were monitored at 2-day old adults.

### Autophagy

Autophagy was measured in hypodermal seam cells of L4 worms according to guidelines (*38*). Autophagosome number was assessed by using the GFP::LGG-1 reporter strain MAH14 grown on OP50 and HT115 bacterial diets for at least two generations. Approximately 20-30 L4-staged animals were collected, anaesthetized with 0.1% sodium azide and mounted on agarose pads for microscopic observation. The number of the GFP::LGG-1 positive autophagic puncta was quantified.

### *rnC* knockout in Nissle 1917

*E. coli* Nissle 1917 *rnc* knock-out strain was created by Red/ET recombination. The protocol was adapted from Datsenko and Wanner (*39*). Wild-type Nissle 1917 strain was cultured in LB overnight at 37°C, 200 rpm. The next day, the kanamycin-resistant pKD4 cassette was amplified by PCR using the following primers 5’-CATCGTAATTAATCGGCTTCAACGGAAGCTGGGCTACACTTGTAGGCTGGAGCTG CTTCG-3’ and 5’-CTGACCTGGCAGTGGATAGTAAATTCCTGATCGTGCGCTTATGGGAATTAGCCATG GTCC-3’. The primers contain overhangs corresponding to the neighboring sequences of the *rnc* gene. In parallel, the pKD46 plasmid was transformed into wild-type Nissle 1917 by electroporation. The next day, the PCR product was transformed into Nissle 1917 by electroporation. Adding 1 mM L-arabinose induced the ʎ recombinase expressed from the pKD46 plasmid, which lead to the exchange of the *rnc* gene with the pKD4 cassette at the corresponding overhangs. Clones of Nissle 1917, where the *rnc* gene was knocked-out and replaced by the kanamycin-resistant pKD4 cassette were picked after kanamycin selection. Knock-out was further confirmed by PCR using the following primers 5’-CTGAAGCGAATCTGGTCGGT-3’ and 5’-CACTTTGTTCACCGCGAGGA-3’.

### Bacterial RNA isolation

A colony of HT115 bacteria were inoculated in LB medium and grown overnight at 37°C in a shaking incubator. Next day 0.5 ml of the overnight culture were inoculated in 10 ml medium containing Tryptone (0.25%), NaCl (0.3%), Cholesterol (5μg/ml), CaCl_2_ (1mM), MgSO_4_ (1mM), KPO_4_ (25mM), Ampicillin (100μg/ml), Nystatin (100U/ml), and grown overnight at 23°C in a shaking incubator. Bacterial pellets were obtained after 3 min centrifugation at 1,500 rcf. RNA was isolated from the pellets using the Quick-RNA Fungal/Bacterial Kit from Zymo Research and stored at -20^0^C till use.

### Total RNAseq

Ribosomal RNA depletion was performed on 300ng *E. coli* total RNA using NEBNext rRNA Depletion Kit Bacteria, (Cat#E7850L, NEB, Ipswich, MA, USA). Following elution in 8µl water, 1µl of eluate for monitoring the depletion of ribosomal RNA on TapeStation instrument (Agilent Technologies, Santa Clara, CA, USA) using the High Sensitivity RNA ScreenTape (Agilent, Cat# 5067-5579), 6µl of eluate were then mixed with Fragment, Prime Finish Mix provided in the TruSeq Stranded Total RNA Library Prep Gold Kit (Cat# 20020599, Illumina, San Diego, CA, USA) used for completing library preparation, in conjunction with the TruSeq RNA UD Indexes (Cat# 20022371, Illumina). 15 cycles of PCR were performed. Libraries were quality-checked on the Fragment Analyzer (Agilent Technologies, Santa Clara, CA, USA) using the Standard Sensitivity NGS Fragment Analysis Kit (Cat# DNF-473, Agilent Technologies) revealing good quality of libraries (average concentration was 72±46 nmol/L and average library size was 317±37 base pairs). Samples were pooled to equal molarity. The pool was quantified by Fluorometry using the QuantiFluor ONE dsDNA System (Cat# E4871, Promega, Madison, WI, USA) and sequenced Single-Reads 76 bases (in addition: 8 bases for index 1 and 8 bases for index 2) on NextSeq 500 using the NextSeq 500 High Output Kit 75-cycles (Illumina, Cat# FC-404-1005). Flow lanes were loaded at 1.8pM. 1% PhiX was included in the pool. Primary data analysis was performed with the Illumina RTA version 2.11.3. This Nextseq runs compiled a large number of reads (on average per sample: 22.5±11.7 millions pass-filter reads).

### RNAseq for small RNAs

The kit QIAseq FastSelect –5S/16S/23S (Cat# 335921, Qiagen, Hilden, Germany) was used for inhibiting the amplification of ribosomal RNA during library preparation which was then performed from 120ng total RNA of E. coli total RNA using SMARTer smRNA-Seq Kit for Illumina (Cat# 635029, Takara Bio, Shiga, Japan). Libraries were quality-checked on the Fragment Analyzer (Agilent Technologies, Santa Clara, CA, USA) using the High Sensitivity NGS Fragment Analysis Kit (Cat# DNF-474, Agilent Technologies) revealing good quality of libraries (average concentration was 0.94±0.22 nmol/L and average library size was 191±5 base pairs). Samples were pooled to equal molarity. The pool was quantified by Fluorometry using the QuantiFluor ONE dsDNA System (Cat# E4871, Promega, Madison, WI, USA) and sequenced Single-Reads 76 bases (in addition: 8 bases for index 1 and 8 bases for index 2) on NextSeq 500 using the NextSeq 500 High Output Kit 75-cycles (Illumina, Cat# FC-404-1005). Flow lanes were loaded at 1.8pM. 1% PhiX was included in the pool. Primary data analysis was performed with the Illumina RTA version 2.11.3. This Nextseq run compiled a large number of reads (on average per sample: 24.0±6.2 millions pass-filter reads).

### RNA-seq data analysis

For the total RNA-Seq, reads were mapped against the Ensembl E. coli K12 DH10B reference genome distributed by iGenomes using STAR v2.7.9 (*40*). Read counts were summarized using the featureCounts function of Subread package v2.0.3. The matrix of uniquely mapped read counts was filtered for features with at least 10 reads in at least 3 samples. Read normalization was computed using the blind variance stabilizing transform as implemented in DESeq2 v1.40.2 (*41*), and used as input for clustering by principal component analysis as implemented by prcomp in R v4.3.0.

For the short read RNA-Seq: Illumina sequencing reads were pre-processed by removing the first three nucleotides and polyA trimming using CutAdapt v3.4 (*42*) as per the manufacturer’s instructions, followed by 3’ quality trimming using Trimmomatic v0.39 (*43*). Adapter trimmed reads were pooled and used as input to the Trinity de novo assembly pipeline v2.11.0 (*44*). The short reads were then aligned against the collection of contigs using Bowtie v1.2.3 (-n 1 -l 10) (*45*). Reads were summarized using featureCounts. 4296 contigs had at least 10 reads in at least 3 samples. Uniquely mapped reads for these features were considered for differential expression using the Wald test as implemented in DESeq2. Significantly differentially expressed genes were considered for differences across the genotype contrasts greater than 2-fold with an adjusted p-value of <=0.01. BLASTn was used to identify differentially expressed contigs which map to the *C. elegans* genome.

### Injections

Total bacterial RNA (85 ng/μl), genomic DNA (70 ng/μl) in water, and water as vehicle control were microinjected directly in the syncytium region of polyQ40-expressing *C. elegans* germlines cultured on OP50 diets. Similarly, 100 ng/μl of each dsRNA generated by *in vitro* transcription was used in a mixture to inject polyQ40-expressing *C. elegans* germlines cultured on OP50 diets. In each case approximately 20 young adult worms were injected. The progeny of injected worms was monitored under the microscope. For each condition about 60 2-day old progeny grown on OP50 were used. Injections were repeated at least 3 times.

### In vitro transcription of bacterial segments

To synthesize dsRNA from the plasmids containing the three bacterial segments that were identified to be differentially expressed between OP50 and OP50(xu363) by the smRNAseq analysis we used the MEGAscript™ T7 Kit (ThermoFisher Scientific). In short we prepared the template DNA by linearizing all three plasmids (digests with HaeII). We obtained shorter regions that contain the DNA segments of interest in between two T7 polymerases. Following the manufacturer’s protocol we assembled the transcription reaction and generated dsRNAs that were subsequently recovered by phenol:chloroform extraction and isopropanol precipitation. The pellets were re-suspended in water and frozen till the day of injection.

### Proteome analysis

Synchronous N2 and AM141 worms were placed on 4 plates and left to lay approximately 500 eggs. Once the worms reached the adult stage were collected in M9 buffer. Floating eggs were removed and the remaining adults were washed with M9 and placed back in NGM plates. The next day, the 2-day old worms were washed 3 times with M9 and the worm pellet was flash frozen in liquid nitrogen and stored at -80^0^C. Worms were resuspended in 5% SDS, 10 mM Tris(2-carboxyethyl)phosphine hydrochloride (TCEP), 0.1 M TEAB and lysed by sonication using a PIXUL Multi-Sample Sonicator (Active Motif) with Pulse set to 50, PRF to 1, Process Time to 20 min and Burst Rate to 20 Hz followed by a 10 min incubation at 95°C. Lysates were TCA precipitated according to a protocol originally from Luis Sanchez (https://www.its.caltech.edu/~bjorker/TCA_ppt_protocol.pdf) as follows. One volume of TCA was added to every 4 volumes of sample, mixed by vortexing, incubated for 10 min at 4°C followed by collection of precipitate by centrifugation for 5 min at 23,000 g. Supernatant was discarded, pellets were washed twice with acetone precooled to -20°C and the washed pellets were incubated open at RT for 1 min to allow residual acetone to evaporate. Pellets were resuspended in 2 M Guanidinium HCl, 0.1 M Ammonium bicarbonate, 5 mM Tris(2-carboxyethyl)phosphine hydrochloride solution (TCEP), phosphatase inhibitors (Sigma P5726&P0044) and proteins were digested as described previously (PMID:27345528). Shortly, proteins were reduced for 60 min at 37°C and alkylated with 10 mM chloroacetamide for 30 min at 37°C. After diluting samples with M Ammonium bicarbonate buffer to a final Guanidinium HCl concentration of 0.4 M, proteins were digested by incubation with sequencing-grade modified trypsin (1/100, w/w; Promega, Madison, Wisconsin) for 12 h at 37°C. After acidification using 5% TFA, peptides were desalted using C18 reverse-phase spin columns (Macrospin, Harvard Apparatus) according to the manufacturer’s instructions, dried under vacuum and stored at -20°C until further use.

Dried peptides were resuspended in 0.1% aqueous formic acid and subjected to LC– MS/MS analysis using a Exploris 480 Mass Spectrometer fitted with an Vanquish Neo (both Thermo Fisher Scientific) and a custom-made column heater set to 60°C. Peptides were resolved using a RP-HPLC column (75 μm × 30 cm) packed in-house with C18 resin (ReproSil-Pur C18–AQ, 1.9 μm resin; Dr. Maisch GmbH) at a flow rate of μL/min. The following gradient was used for peptide separation: from 4% B to 10% B over 5 min to 35% B over 45 min to 50% B over 10 min to 95% B over 1 min followed by 10 min at 95% B to 5% B over 1 min followed by 4 min at 5% B. Buffer A was 0.1% formic acid in water and buffer B was 80% acetonitrile, 0.1% formic acid in water.

The mass spectrometer was operated in DIA mode with a cycle time of 3 seconds. MS1 scans were acquired in the Orbitrap in centroid mode at a resolution of 120,000 FWHM (at 200 m/z), a scan range from 390 to 910 m/z, normalized AGC target set to 300 % and maximum ion injection time mode set to Auto. MS2 scans were acquired in the Orbitrap in centroid mode at a resolution of 15,000 FWHM (at 200 m/z), precursor mass range of 400 to 900, quadrupole isolation window of 12 m/z with 1 m/z window overlap, a defined first mass of 120 m/z, normalized AGC target set to 3000% and a maximum injection time of 22 ms. Peptides were fragmented by HCD (Higher-energy collisional dissociation) with collision energy set to 28% and one microscan was acquired for each spectrum.

The acquired raw-files were searched using the Spectronaut (Biognosys v17.4) directDIA workflow against a *C. elegans* database (consisting of 26585 protein sequences downloaded from Uniprot on 20220222) and 392 commonly observed contaminants. Quantitative data was exported from Spectronaut and analyzed using the MSstats R package v.4.7.3. (https://doi.org/10.1093/bioinformatics/btu305).

### GO term analysis and functional association networks

GO term analysis and protein-protein interaction networks enrichments analysis were performed with the use of ShinyGO (ver. 0.77) (http://bioinformatics.sdstate.edu/go/) with 0.01 FDR cutoff and STRING (ver. 12.0) (https://string-db.org/cgi/input?sessionId=b0IlBAPfUcQa&input_page_show_search=on). A q-value of less than 0.05 was used to filter significant changes prior to the pathway analyses. Proteins between -0.3 and +0.3 fold change (log2ratio) were excluded from the analysis.

### Statistical analysis

Statistical analyses and graphs were prepared using the Prism software package (version 9; GraphPad Software; https://www.graphpad.com). Data are reported as the mean values ± standard deviation (SD). For statistical analyses, p values were calculated by unpaired Student’s t-test and one-way ANOVA with multiple comparisons test. The significance was determined by the p-values: * p < 0.05, ** p < 0.01, *** p < 0.001 and n.s. = not significant p > 0.05.

## Acknowledgments

We thank Pascal Ankli, Jessica Arnegger, Tara Thorsen, Vanessa Brullo, Sheuli Begum, Jordi Greoles Cano, Isabella Santi, Louise Larsson, Médéric Diard and Dora Stetak for technical support. We also thank the Genomics Facility Basel of the University of Basel and the Department of Biosystems Science and Engineering, ETH Zurich for carrying out the Next-Generation Sequencing. We are grateful to Susan E. Mango, Ian G. Macara and Nikolaos Charmpilas for critical reading of the manuscript. Some nematode strains used in this work were provided by the Caenorhabditis Genetics Center, which is funded by the National Center for Research Resources of the National Institutes of Health and S. Mitani (National Bioresource Project) in Japan. We thank Read Pukkila-Worley and Andy Fire for providing the AU306 strain and plasmid vectors, respectively.

## Funding

This work is supported by

Swiss National Science Foundation grant 185127 (AS)

Swiss National Science Foundation grant 197779 (AS)

The Novartis Foundation for Medical-Biological Research grant #20C179 (AS)

University of Basel grant FoFo 3BZ5106 (AS)

University of Basel

## Author contributions

Conceptualization: EK, AS

Methodology: EK, CM, GF

Investigation: EK, CM, GF, DR, AS

Visualization: EK, CM, GF

Funding acquisition: EK, AS

Supervision: EK, AS

Writing – original draft: EK, AS

Writing – review & editing: EK, AS

## Competing interests

Authors declare no competing interests.

## Notes

### Competing Interest Statement

The authors have declared no competing interest.

